# Non-classical glycosylation determines intracellular trafficking of APP and Aβ production

**DOI:** 10.1101/2022.06.20.496769

**Authors:** Yuriko Tachida, Junko Iijima, Kazuto Takahashi, Hideaki Suzuki, Yoshiki Yamaguchi, Katsunori Tanaka, Miyako Nakano, Daisuke Takakura, Nana Kawasaki, Yuko Saito, Hiroshi Manya, Tamao Endo, Shinobu Kitazume

**Affiliations:** Disease Glycomics Team, RIKEN, 2-1 Hirosawa, Wako 351-0198, Saitama, Japan; Department of Clinical Laboratory Sciences, School of Health Sciences, Fukushima Medical University School of Medicine, Fukushima 960-8516, Japan; Division of Pharmaceutical Physical Chemistry, Tohoku Medical and Pharmaceutical University, Miyagi 981-8558, Japan; Department of Chemical Science and Engineering, School of Materials and Chemical Technology, Tokyo Institute of Technology, 2-12-1 Ookayama, Meguro-ku, Tokyo 152-8550, Japan; Biofunctional Synthetic Chemistry Laboratory, RIKEN Cluster for Pioneering Research, 2-1 Hirosawa, Wako, Saitama 351-0198, Japan; Graduate School of Integrated Sciences for Life, Hiroshima University, Higashi-hiroshima 739-8530, Japan; Graduate School of Medical Life Science, Yokohama City University, Yokohama 230-0045 Japan; Department of Neuropathology, Tokyo Metropolitan Geriatric Hospital and Institute of Gerontology, Tokyo 173-0015, Japan; Molecular Glycobiology, Research Team for Mechanism of Aging, Tokyo Metropolitan Geriatric Hospital and Institute of Gerontology, Tokyo 173-0015, Japan

**Keywords:** Alzheimer’s disease (AD), Amyloid β (Aβ), Amyloid precursor protein (APP), intracellular trafficking, O-glycosylation

## Abstract

A primary pathology of Alzheimer’s disease (AD) is Aβ deposition in brain parenchyma and blood vessels, the latter being called cerebral amyloid angiopathy (CAA). Parenchymal amyloid plaques presumably originate from neuronal Aβ precursor protein (APP), but vascular amyloid deposits’ origins remain unclear. Endothelial APP expression in APP-knock-in mice was recently shown to expand CAA pathology, highlighting endothelial APP’s importance. Furthermore, two types of endothelial APP—with and without O-glycans—have been biochemically identified, but only the former is cleaved for Aβ production, indicating the critical relationship between APP O-glycosylation and processing. Here, we analyzed APP glycosylation and its intracellular trafficking in neurons and endothelial cells. Although protein glycosylation is generally believed to precede cell surface trafficking, which was true for neuronal APP, we unexpectedly observed that APP lacking O-glycans is externalized to the endothelial cell surface and transported back to the Golgi apparatus, where it then acquires O-glycans. Knockdown of genes encoding enzymes initiating APP O-glycosylation significantly reduced Αβ production, suggesting this non-classical glycosylation pathway contributes to CAA pathology and is a novel therapeutic target.

## Introduction

Alzheimer’s disease (AD) is a progressive neurodegenerative disorder that features two pathological hallmarks: intraneuronal neurofibrillary tangles^1^ and extracellular deposition of amyloid β (Aβ)^2^. Aβ is generated from amyloid precursor protein (APP). When APP is cleaved at the plasma membrane at the α-site within the Aβ sequence, N-terminal ectodomain, sAPPα, is released, and subsequent γ-secretase cleavage of the carboxy-terminal fragment generates p3 peptide instead of Aβ^3^. While part of cell surface APP is internalized and transported to endosome, BACE1 protease cleaves at the β-site during the endocytic pathway^4^, leading to shedding of the N-terminal ectodomain, sAPPβ, and subsequent cleavage of the carboxy-terminal fragment at the γ-site to produce Aβ. Therefore, the cellular level of Aβ production largely depends on the extent to which APP encounters each active secretase in the cell^5^.

Both APP and its secretase are glycosylated, and several reports suggest that such glycosylation affects Aβ production. Unusual GalNAc-type O-glycosylation to a Tyr residue within the Aβ sequence, which is frequently found in cerebrospinal fluid from AD patients^6^, results in conformational changes of APP favorable for the amyloidogenic pathway^7^. Modification of the N-glycans of BACE1 with bisecting GlcNAc attenuates its lysosomal targeting and enhances Aβ production^8^, indicating that glycosylation can affect the intracellular localization of secretases to modulate Aβ production. APP has two N-glycans at specific Asn residues and multiple GalNAc-type O-glycans at Ser/Thr residues^9–12^. N-glycosylation is initiated in the ER to generate oligomannose-type glycan; then, a series of Golgi-resident glycosidases and glycosyltransferases functions in the processing of N-glycans and the addition of GalNAc-type O-glycans^13^, which are considered to be maturation steps of glycoproteins and necessary for their trafficking to functional locations. Notably, O-GalNAc glycoproteome analysis revealed that remarkable numbers of Golgi- and ER-resident proteins have O-glycans^9^.

Aβ plaques are not limited to brain parenchyma and vascular Aβ deposition, known as cerebral amyloid angiopathy (CAA), is also observed at a high frequency^14^. Parenchymal Aβ is considered to originate from neuronal APP, but a recent finding showing that endothelial APP expression^15^ contributes to vascular Aβ deposition^16^ highlighted the importance of endothelial APP for the pathogenesis of CAA. Cell-type-specific mRNA splicing produces different APP isoforms, namely, neuronal APP695 and endothelial APP770, with APP770 having additional OX2 and KPI domains compared with APP695. In this study, we focused on the glycosylation and intracellular trafficking of neuronal and endothelial APP. Contrary to neuronal APP, in which both N- and O-linked glycans are attached to the APP before its cell surface transport, we found that endothelial APP takes a non-classical biosynthetic pathway; APP lacking O-glycans, but having N-glycans, is transported to the cell surface, and is then internalized and transported back to the Golgi apparatus for O-glycosylation. Our study sheds light on an overlooked functional connection between cell-type-dependent protein glycosylation and intracellular trafficking, and also raises the possibility that modulation of the O-glycosylation pathway could attenuate cellular Aβ production.

## Results

### O-glycosylated sAPP is shed from neurons and endothelial cells

Only limited information is available concerning APP O-glycosylation sites, and therefore we first conducted site-specific mapping of APP O-glycans using a mass spectrometry-based method. Hemagglutinin (HA)-tagged human APP770 was expressed in HEK293T cells, and HA-sAPP770 purified from culture medium was treated with trypsin plus OpeRATOR protease^17^, and used for MS/MS analysis. In addition to several known O-glycosylation sites^9, 10^, we additionally identified Thr269, Thr274, Thr352, Thr366, Thr367, Ser370, Thr381, Thr651, Thr652, and Thr663 as novel O-glycosylation sites (Fig 1A and B, Fig EV1). Notably, O-glycosylation sites of APP are concentrated at two sites: one near the KPI plus OX2 domain, and the other near the β-cleavage sites.

**Figure 1.**
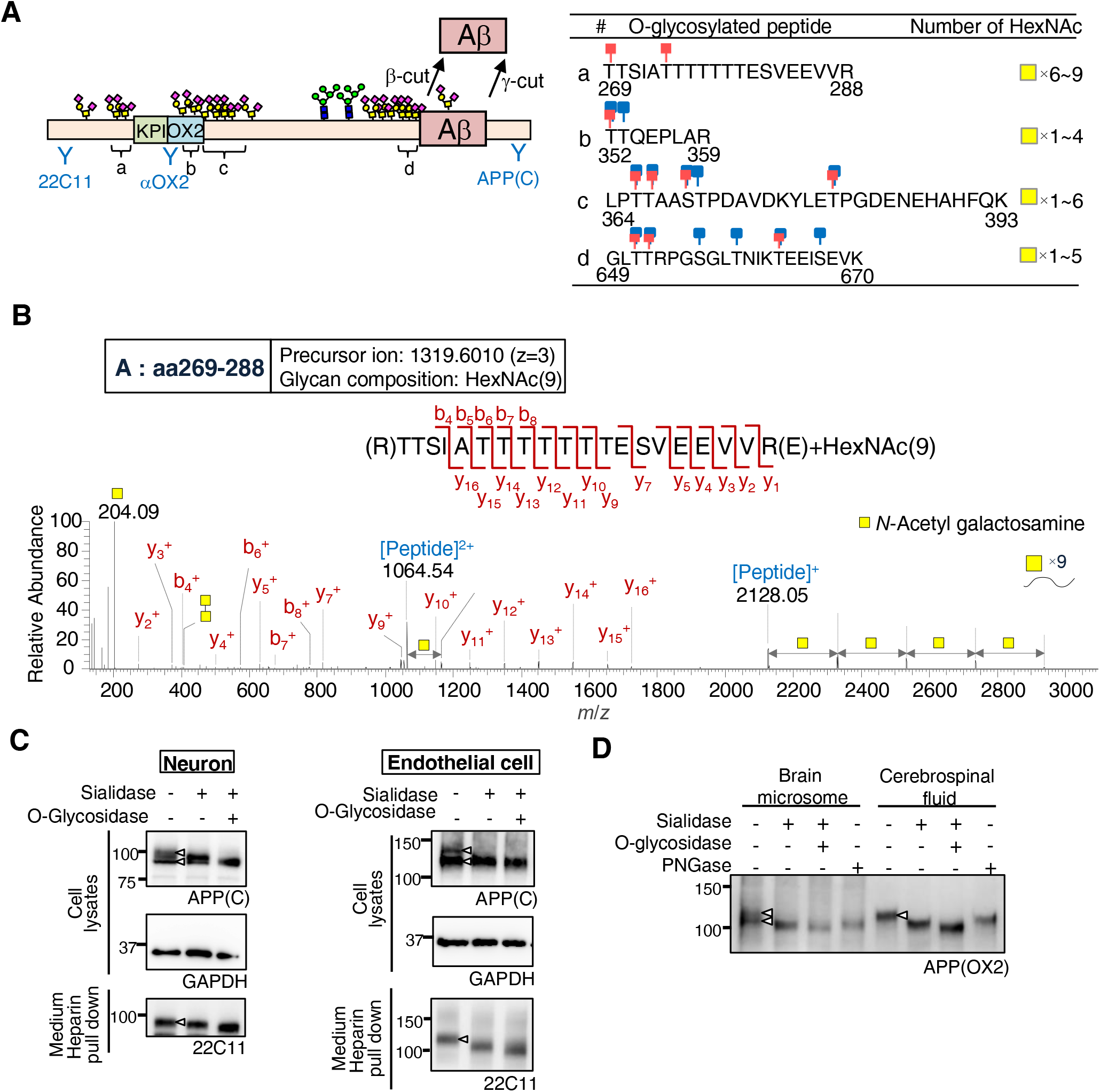
O-glycosylated APP is preferentially secreted. A Schematic of APP770 structure, in which N- and O-glycosylation sites and site-specific anti-APP antibodies are shown. The positions of identified O-glycosylation peptides are also shown (a–d). The table shows peptide sequences with known O-glycosylation sites (blue) and newly identified O-glycosylation sites (red), and a number of identified HexNAc residues are also shown. B Product ion spectrum of APP770 O-glycopeptide (aa269–288) arising from the precursor ion at *m/z* 1319.6010 (z=3). C Lysates and sAPP pulled down with heparin agarose from the medium of mouse primary neurons and endothelial cells were treated with sialidase or O-glycosidase, and analyzed by immunoblotting for APP and GAPDH (loading control). D Human brain microsomes and cerebrospinal fluid were incubated with heparin agarose to pull down APP and sAPP, respectively. The samples were treated with sialidase, O-glycosidase, or PNGase, and blotted for APP770 and sAPP770, respectively.

We then analyzed neuronal and endothelial APP, and western blot analysis using anti-C-terminal APP antibody revealed that full-length APP exhibits double bands in both types of cell lysate. After treatment with sialidase plus O-glycosidase, the latter being an enzyme that specifically removes non-sialylated core 1 O-glycan disaccharide (Galβ1-3GalNAcα1-Ser/Thr), in both cell types the upper APP band disappeared and merged with the lower band (Fig 1C), indicating that the upper band represented core 1-type O-glycosylated APP and the lower band represented APP lacking O-glycans. Meanwhile, sAPP, released into the medium from both types of cell, was most sensitive to sialidase plus O-glycosidase, indicating that sAPP was mostly O-glycosylated. We then extended this analysis to human brain samples. Brains contain a mixture of different APP isoforms (APP695, 751, and 770), making analysis difficult. We therefore used anti-OX2 antibody and focused on APP770, which is heavily O-glycosylated and thus easily differentiated from its non-O-glycosylated form by SDS-PAGE. Again, the human brain microsome fractions contained both forms of APP770, whereas the sAPP770 in the cerebrospinal fluid (CSF) was fully O-glycosylated (Fig 1D).

### Endothelial cell surface APP receives O-glycan sialylation

It was considered that the unique feature of APP of O-glycosylated APP and non-O-glycosylated APP being clearly separated by SDS-PAGE would provide us with a unique opportunity to clarify the role of the O-glycosylation pathway in intracellular APP trafficking. We first performed a cell surface biotinylation experiment using neurons and endothelial cells. In neurons, only the upper band was biotinylated, indicating that APP was transported to the cell surface after O-glycosylation (Fig 2A). Unexpectedly, however, in endothelial cells not only O-glycosylated but also non-O-glycosylated APP was biotinylated. These results indicate that, in neurons, fully glycosylated APP is selectively transported to the cell surface, while in endothelial cells, APP can be transported to the cell surface without O-glycan. The latter result contradicts the accepted belief that nascent proteins receive O-glycans in the Golgi apparatus before reaching the cell surface^18^, and does not fully explain why sAPP is exclusively O-glycosylated. One possibility, namely, that non-O-glycosylated APP could be less stable than O-glycosylated APP and only the latter survives, was ruled out by half-life analysis, as the half-life of cell surface biotinylated non-O-glycosylated APP was ∼21 h, which was ∼5 times longer than that of O-glycosylated APP (4.1 h) (Fig 2B).

**Figure 2.**
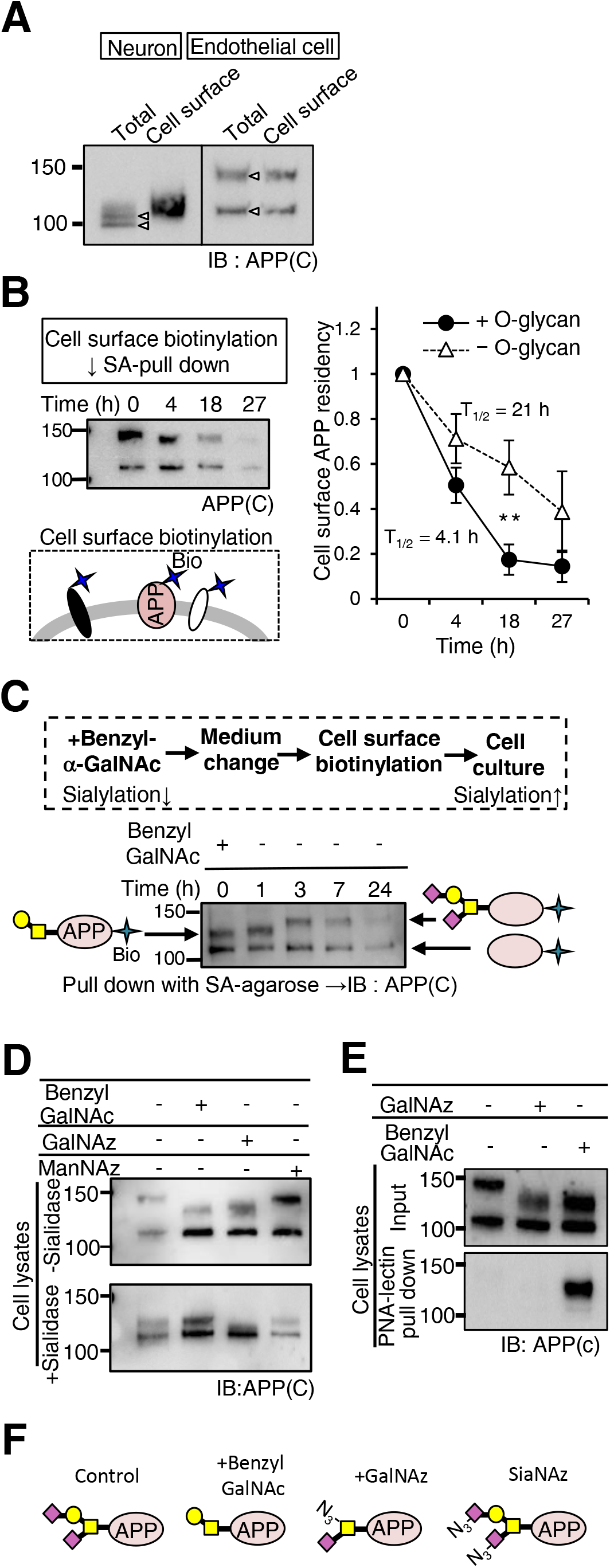
Cell surface APP is internalized and its O-glycans are sialylated. A After cell surface labeling with NHS-LC-biotin and subsequent incubation of mouse primary neurons and human brain microvascular endothelial cells (BMECs), biotinylated proteins were pulled down with streptavidin (SA)-agarose and blotted for APP. B After cell surface labeling with sulfo-NHS-LC-biotin and subsequent incubation, biotinylated proteins were pulled down with SA-agarose and blotted for APP. On the basis of the quantification of biotinylated APP, the half-life of cell surface APP was expressed as mean ± SEM (n=3 independent western blots). **p < 0.01, Student’s t-test. C After BMECs were incubated with benzyl-α-GalNAc for 16 h, the cells were surface-biotinylated and cultured again in the absence of benzyl-α-GalNAc for different periods. Biotinylated proteins pulled down with SA-agarose were blotted for APP. D BMECs were cultured with benzyl GalNAc, Ac_4_GalNAz, or Ac_4_MaNAz, and the cell lysates were treated with sialidase and blotted for APP. The upper band of all samples was sensitive to sialidase, indicating that GalNAz incorporation allowed sialylation. E BMECs were cultured with benzyl-α-GalNAc or Ac_4_GalNAz. The cell lysates were incubated with immobilized PNA lectin that detects the Galβ1,3GalNAc structure. PNA lectin-precipitated samples were then blotted for APP. GalNAz-incorporated APP was not reactive with PNA lectin, indicating impaired galactosylation. F Considering the results in (D) and (E), typical O-glycan structure of APP, treated with benzyl-α-GalNAc, GalNAz, or ManAz (for SiaNAz), is shown.

Another possibility is that cell surface non-O-glycosylated APP is internalized and undergoes O-glycosylation within the Golgi apparatus before processing. To explore this possibility, we designed an experiment that used a combination of cell surface biotinylation and treatment of cells with benzyl-α-GalNAc, a drug that perturbs the extension of O-glycans, such as the sialylation of core 1-type O-glycan^19^. After treatment, the benzyl-α-GalNAc was removed from the medium and the cells were labeled with biotin. Cell surface biotinylated proteins were enriched by streptavidin (SA)-agarose. We first checked that the molecular weight of biotinylated O-glycosylated APP was reduced by this treatment (Fig 2C)^12^. Indeed, the benzyl-α-GalNAc-treated APP was hyposialylated (Fig 2D), but had Galβ1,3GalNAc residues, based on the reactivity to PNA lectin (Fig 2E and F). Interestingly, further incubation in the absence of benzyl-α-GalNAc led to an increase in the molecular weight of biotinylated O-glycosylated APP. These results indicate that cell surface APP is internalized and then modified with sialic acid to produce extended O-glycans.

### Non-O-glycosylated APP at the endothelial cell surface is internalized and receives O-glycans

We then hypothesized that the initial O-GalNAc transfer event might also occur upon the internalization of cell surface APP. Such GalNAc residues can be metabolically labeled using a GalNAc analog, GalNAc-azide (Ac_4_GalNAz), which is incorporated into O-glycans via endogenous biosynthetic pathways and can be covalently tagged with an azide-reactive probe^20, 21^. Thus, we performed cell surface biotinylation and subsequent O-glycan metabolic labeling with Ac_4_GalNAz. Using an adenovirus technique, APP-FLAG was expressed in endothelial cells, which were treated with a peracetylated derivative, either Ac_4_GalNAz or Ac_4_ManNAz, to visualize GalNAc or sialic acid, respectively^22, 23^. The incorporated azide sugars (GalNAz or SiaNAz) were click-labeled with TAMRA-conjugated dibenzocyclooctyne (DIBO)^24^. We detected fluorescent signals corresponding to the O-GalNAz glycosylated APP-FLAG, verifying the incorporation of GalNAz into APP (Fig 3A). The incorporation of GalNAz into APP was also examined by mass spectrometry analysis of the immunopurified APP-derived glycopeptide (Appendix Fig S1). Furthermore, analysis with immunofluorescence microscopy revealed that most of the intracellular APP signals co-localized with GalNAz signals (Fig 3B). Notably, part of GalNAz is enzymatically converted to UDP-GlcNAz in addition to UDP-GalNAz and nuclear O-GlcNAcylated proteins are presumably O-GlcNAz-labeled^25^. Indeed, in the mutant CHO cell line, IdlD cells, which lack UDP-galactose epimerase (GALE) activity and are unable to convert GalNAz to GlcNAz, we found no nuclear azide signal, with all of the signals instead being found in the intracellular vesicles (Fig EV2)^25^. Next, O-GalNAz glycans as well as APP were visualized with several organelle markers in endothelial cells. As has been reported in other cells^26, 27^, APP co-localized with the trans-Golgi marker adaptin-γ, the recycling endosome marker Rab11, the early endosome markers EEA1 and Rab5, and, to a lesser extent, the late endosome marker Rab7 (Fig EV3A and B). In comparison with APP, the O-GalNAz glycan signal co-localized less with adaptin-γ and EEA1 but was relatively enriched in the recycling endosome (Fig EV3C and D). Next, we combined the cell surface biotinylation experiment with metabolic sugar labeling. After cell surface biotinylation and subsequent metabolic labeling with Ac_4_GalNAz or Ac_4_ManNAz, we found that each biotinylated APP had a fluorescent signal derived from azide sugars (Fig 3C), clearly demonstrating that cell surface biotinylated APP was internalized and then underwent O-GalNAz glycan modification. Notably, the molecular weight of GalNAz-labeled APP was lower than that of wild-type APP. On the basis of the analysis of GalNAz-labeled APP with sialidase and PNA lectin, we found that GalNAz incorporation allowed sialylation but blocked subsequent galactosylation (Fig 2D, E and F). However, sAPP was still generated and fluorescently detected in the culture medium (Fig 3A), indicating that the Ac_4_GalNAz treatment did not impair the overall intracellular trafficking of APP.

**Figure 3.**
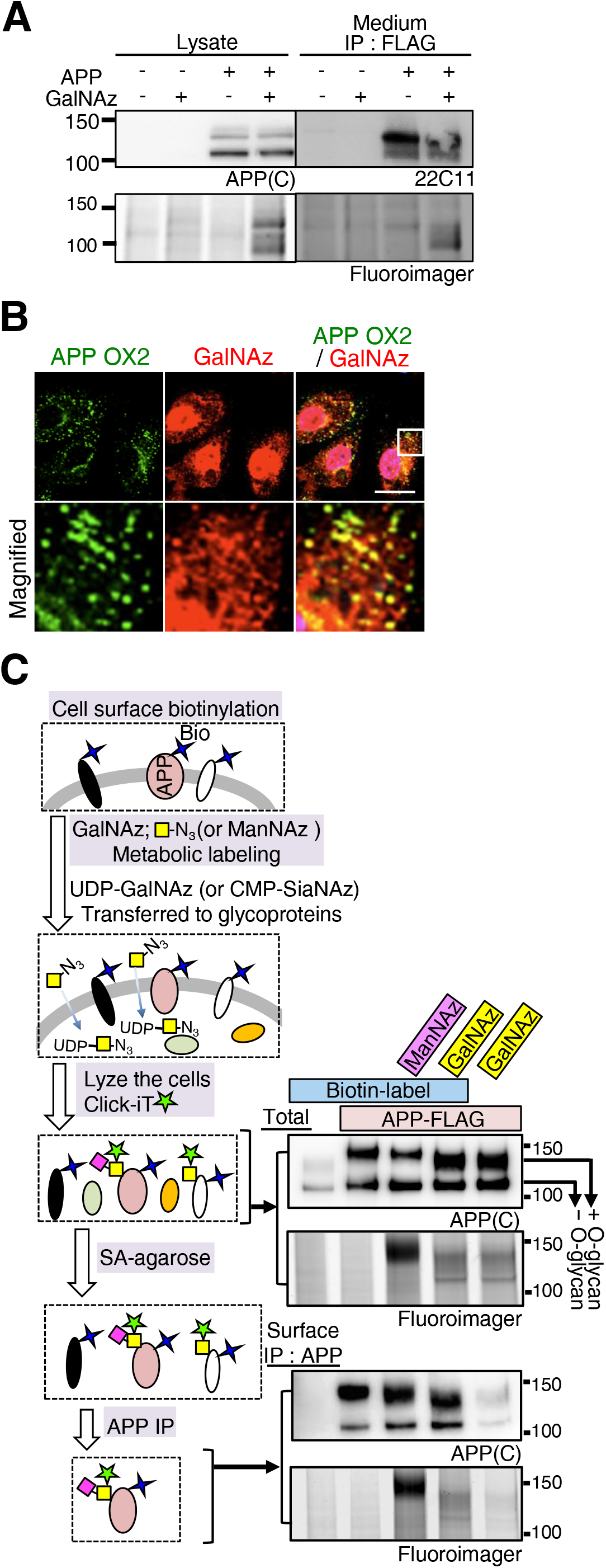
Cell surface APP is internalized and O-glycosylated. A BMECs expressing FLAG-APP with an adenovirus system were metabolically labeled with GalNAz. The cell lysates were treated with TAMRA-conjugated DIBO, and sAPP pulled down with anti-FLAG (M2)-coupled agarose was blotted for APP or analyzed by a fluoroscanner. B Immunofluorescence microscopy shows that endothelial APP co-localizes with O-GalNAz glycan signals. Scale bar, 20 μm. Lower panels show magnified images. C After cell surface biotinylation, cells were cultured with Ac_4_GalNAz or Ac_4_ManNAz for 6 h, and the lysates were then treated with TAMRA-conjugated DIBO to fluorescently label the azide group. The presence of O-GalNAz glycan in the biotinylated APP was assessed by a fluoroscanner.

### APP O-glycosylation affects Aβ generation

Even though APP is transported to the cell surface in an O-glycosylation-independent manner, sAPP is exclusively O-glycosylated, raising the possibility that the level of O-glycan in APP could affect its processing. Twenty polypeptide GalNAc-T (GalNAc-T) genes have been identified as being involved in the initiation of O-glycosylation of proteins in humans^18^. First, to identify the responsible GalNAc-T enzyme(s), three kinds of APP-derived peptide encoding O-glycosylation sites were incubated with a series of recombinant GalNAc-Ts and UDP-GalNAc *in vitro*, and the reaction products were analyzed by mass spectrometry^28^. In addition to GalNAc-T2 and -T3, which have been previously reported to transfer GalNAc to APP^9^, we found that GalNAc-T6, which has the highest homology with GalNAc-T3 (Appendix Fig S2A)^18^, could also transfer GalNAc to all of the peptides (Fig. 4A, Appendix Fig S2B)^18^. Other GalNAc-Ts exhibited negligible activity. Notably, the relative GalNAc-T6 activities for all three peptides were higher than those of GalNAc-T2 and -T3, and we could even detect the product in which both of the two O-glycosylation sites (Thr366 and Thr367) were occupied by GalNAc. To date, we have analyzed 10 of 20 GalNAc-Ts. However, GalNAc-T4, -T12, and -T14, which have higher homology with GalNAc-T2, -T3, and - T6, showed no activity with any of the peptide substrates, suggesting that GalNAc-T2, - T3, and -T6 are the major APP O-glycosylation enzymes.

**Figure 4.**
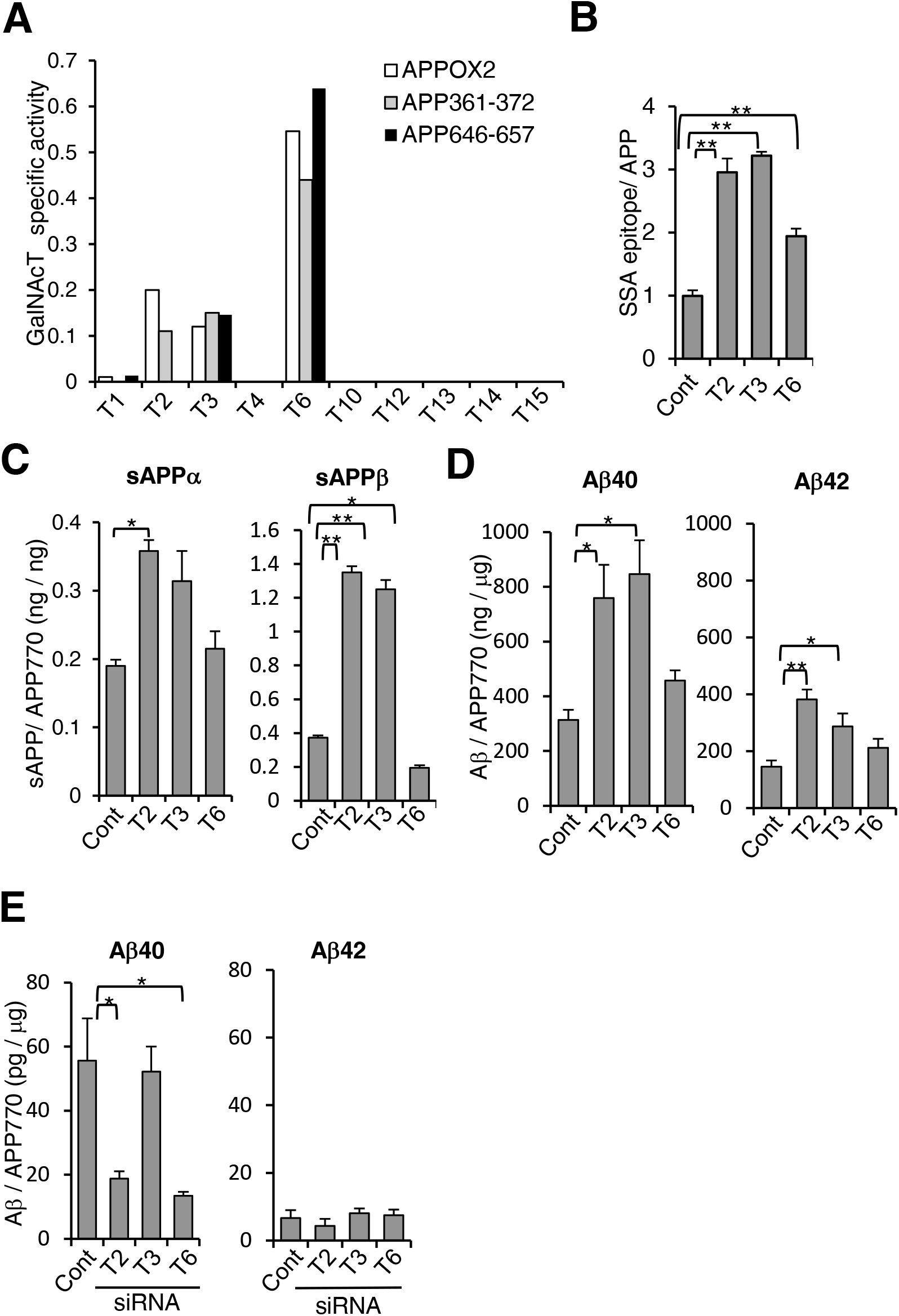
Modulation of the APP O-glycosylation enzyme affects Aβ generation. A Three kinds of APP-derived peptides were incubated with a series of GalNAc-T enzymes and UDP-GalNAc, and the reaction products were analyzed by MS. The specific activity of GalNAc-Ts was calculated from the ratio of signal intensity of the GalNAc-incorporated peptide to that of the acceptor peptide. B Lysates of BMECs transfected with GalNAc-T2, -T3, -T6, or control vector were analyzed by SSA lectin ELISA and APP levels to measure O-glycans on APP. Data show mean ± SEM, n=3. C,D BMECs were transfected with GalNAc-T2, -T3, -T6, or control vector. The levels of sAPPα and sAPPβ in the medium (C) or intracellular Aβ40/42 (D) were measured and are shown as the mean ± SEM, n=6. E HeLa cells were transfected with GalNAc-T2, T3, and T6 or control siRNA. The levels of intracellular Aβ40 and 42 were measured and are shown as mean ± SEM. Statistical analysis was mainly performed by one-way ANOVA with Dunnett’s multiple comparison test or Tukey-Kramer test (for d, Aβ42); *p < 0.05, **p < 0.01.

To define whether or not these GalNAc-Ts act on APP O-glycosylation in the cell, we developed an APP sandwich lectin ELISA system with *Sambucus sieboldiana* agglutinin (SSA) that detects α2,6-sialylated O-GalNAc glycan. The overexpression of GalNAc-T2, -T3, and -T6 markedly increased the level of sialylated O-glycan on APP (Fig 4B). We then investigated the effect of APP O-glycosylation on APP processing and Aβ production. Overexpression of GalNAc-T2 and -T3 resulted in significant increases in sAPPα (∼2-fold), sAPPβ (∼3-fold) (Fig 4C), and Aβ40/42 (∼2-fold) (Fig 4D). GalNAc-T6 exhibited higher enzyme activity to APP *in vitro*, whereas overexpression of GalNAc-T6 did not increase the secretion of sAPP. Previous reports showed that GalNAc-T6 overexpression reduces the level of E-cadherin^29^, and a reduction in this cell adhesion molecule could affect intracellular APP sorting. Partial knockdown of GalNAc-T2 and -T6, but not -T3, in HeLa cells significantly reduced the cellular level of Aβ40 but not Aβ42 (Fig 4E). Taken together, these findings indicate that APP O-glycosylation regulates APP processing.

### Internalized APP encounters O-GalNAc enzymes in the Golgi apparatus

Immunofluorescence microscopic analysis confirmed that GalNAc-T2 mostly co-localized with the trans-Golgi marker adaptin-γ^11, 18^, and overlapped with APP and O-GalNAz signals (Fig EV4A). We therefore expected that the internalized APP would be transported to the Golgi apparatus for its O-glycan modification. To investigate this, Halo-tagged APP was used to specifically label cell surface APP. We first prepared two types of Halo-tagged APP, Halo-APP, in which the N-terminal extracellular domain was tagged with Halo, and APP-Halo, in which the C-terminal cytoplasmic region was tagged with Halo, and determined that both could be labeled with membrane-permeable HaloTag TAMRA ligand (Fig EV4B). In contrast, only Halo-APP was labeled with the non-permeable HaloTag Alexa488 ligand, indicating the feasibility of analyzing the fate of cell surface APP after internalization.

During biochemical analysis of internalized Halo-APP, we unexpectedly found that non-permeable HaloTag Alexa488 and biotin ligands exhibited different binding activity to internalized Halo-APP. The HaloTag biotin ligand bound to both O-glycosylated and non-O-glycosylated APP (Fig 5A), whereas the HaloTag Alexa488 ligand bound almost exclusively to the O-glycosylated APP (Fig 5B). These results indicated that, using these ligands, internalized O-glycosylated APP (Alexa488-Halo ligand) could be discriminated from internalized APP overall (Biotin-Halo ligand). Notably, the internalized APP significantly co-localized with GalNAc-T2 (Fig 5C and D) and with T-synthase, a core 1 O-glycan galactosyltransferase (Fig EV4C)^30^, while internalized O-glycosylated APP exhibited poor co-localization with GalNAc-T2 enzyme (Fig. 5D and E). These results indicate that internalized non-O-glycosylated APP is transported to the Golgi for O-glycosylation, and after which it leaves the Golgi and is transported to different locations. To further dissect the intracellular trafficking of internalized O-glycosylated APP, human endothelial cells expressing Halo-APP were incubated with HaloTag Alexa488 ligand for different periods, fixed, and analyzed. As compared with the early internalization process (5 min), further incubation (30 min) led to more co-localization of the internalized O-glycosylated with Rab11 (Fig. EV5A and B), suggesting that one of the destinations to which O-glycosylated APP leaving the Golgi is transported is a recycling endosome.

**Figure 5.**
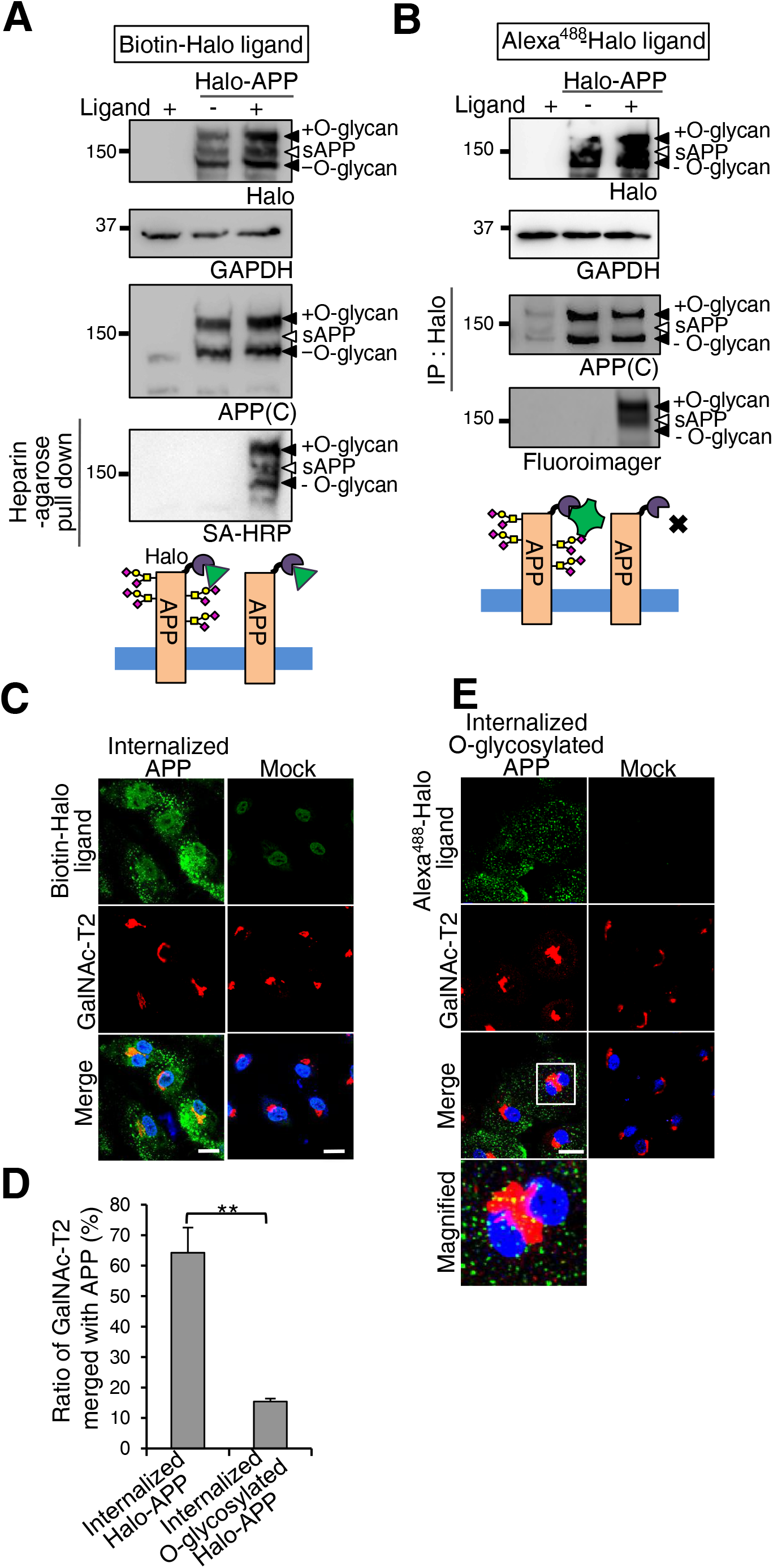
Internalized non-O-glycosylated APP is transported to the Golgi apparatus to encounter GalNAc-T2. A-E BMECs expressing Halo-APP with an adenovirus system were treated with HaloTag biotin ligand and Alexa488-labeled SA (A, C) or HaloTag Alexa488 ligand (B, E). The cell lysates and sAPP pulled down with heparin-Sepharose from the culture medium were analyzed by western blot for APP or HaloTag ligand-coupled APP (A, B). For immunofluorescence, the cells were fixed and immunostained for GalNAc-T2 (C, E). Scale bar, 20 μm. (D) On the basis of the images shown in (C) and (E), the area of co-localization of GalNAc-T2 with internalized Halo-APP was determined as mean ± SEM. **p < 0.01, Student’s t-test.

## Discussion

In protein glycosylation, it is generally believed that newly synthesized proteins undergo both N- and O-glycosylation before being delivered to the cell surface. However, several recent reports show exceptions to this classical glycosylation pathway; extrinsic sialylation, in which IgG and other serum glycoproteins are sialylated by serum-localized nucleotide sugar donor CMP-sialic acid, has been reported by several groups^31, 32^. In this study, we demonstrated another non-classical glycosylation pathway that regulates intracellular APP trafficking in endothelial cells. We found that, in neurons, both N- and O-glycosylated APP are transferred to the cell surface, while in endothelial cells, APP without O-glycans arrives at the cell surface but is then internalized for retrograde transport to the Golgi apparatus where it undergoes O-glycosylation (Fig 6). By using two kinds of HaloTag ligand, we coincidentally succeeded in differentiating the internalization of O-glycosylated APP from that of non-O-glycosylated APP. However, the reason why the bulky and hydrophobic HaloTag Alexa488 ligand binds exclusively to the O-glycosylated Halo-APP remains unclear. We found that the internalized APP significantly co-localized with O-glycosylation enzymes in the Golgi, while internalized O-glycosylated APPs exhibited markedly less co-localization to these O-glycosylation enzymes. Moreover, immunofluorescent microscopy showed that, compared with the intracellular APP signal, the O-GalNAz glycan signal revealed less Golgi localization. These findings suggest that internalized non-O-glycosylated APP is transported in a retrograde fashion to meet the Golgi O-glycosylation enzymes, and fully O-glycosylated APP leaves the Golgi and is transferred differently for processing. An impaired endocytic pathway is implicated in AD, and several molecules, such as low-density-lipoprotein receptor (LDLR) family proteins and PICALM, are reported to be possible sorting receptors for APP and Aβ^5, 33^. One interesting possibility is that these sorting receptors recognize the O-glycosylation status of APP via as-yet-undefined glycan recognition molecules.

**Figure 6.**
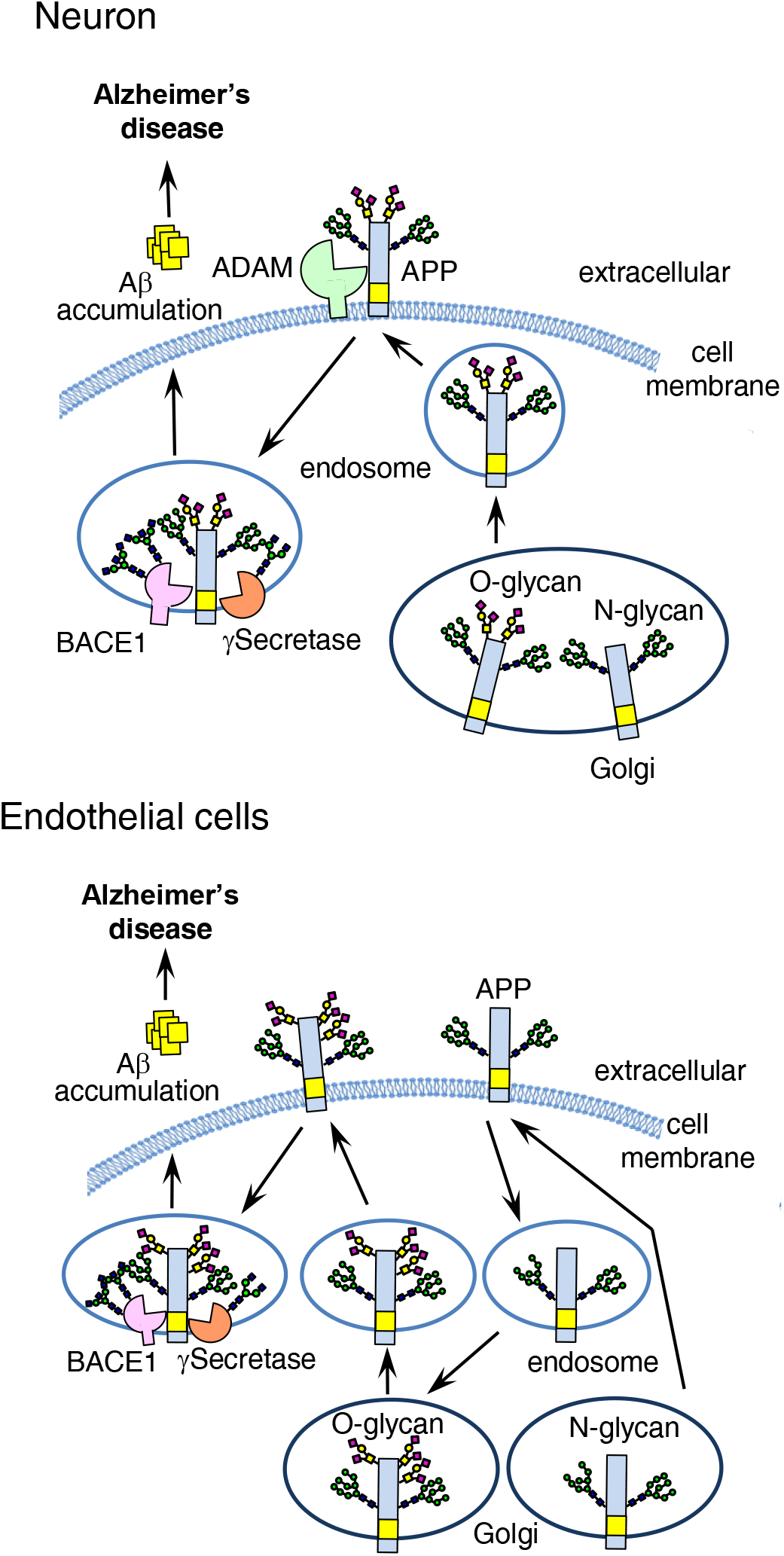
Intracellular trafficking and glycosylation of APP differ among cell types. In neurons, both N- and O-glycosylated APP are transferred to the cell surface, then internalized APP undergoes processing by BACE 1 and γ-secretase to generate Aβ. In the endothelial cells, N-glycosylated APP is transported to the cell surface regardless of O-glycosylation. Non-O-glycosylated APP is internalized and transported in a retrograde fashion to the Golgi apparatus where it undergoes O-glycosylation. The O-glycosylated APP exhibits decreased co-localization with Golgi-resident O-glycosylation enzymes and is eventually processed to generate Aβ.

Although our *in vitro* and cell-based analyses show that GalNAc-T2 is the enzyme responsible for endothelial APP O-glycosylation, GalNAc-T2 siRNA led to a partial reduction in Aβ. Compensatory GalNAc-T3 and T6 activities could contribute to this phenomenon. Furthermore, as several physiological substrates, such as apoC-III and LRP1, have been reported for GalNAc-T2^11^, it is unclear whether GalNAc-T2 inhibition could disturb lipid metabolism and vesicle sorting. Although GalNAc-T6 exhibited higher enzyme activity to APP *in vitro*, the overexpression of GalNAc-T6 did not increase the secretion of sAPP. A previous study has reported that GalNAc-T6 overexpression reduces the level of E-cadherin^29^, and a reduction in such a cell adhesion molecule could also affect the intracellular sorting machinery.

As another important aspect of APP O-glycosylation, we could only observe an effect of GalNAc-T overexpression on the production of sAPP770 and Aβ in cells cultured in low-glucose medium (∼5 mM), in which the level of UDP-GalNAc was limited^34^. Given that UDP-GlcNAc is the end product of the hexosamine pathway and GALE effectively converts UDP-GlcNAc to UDP-GalNAc^35^, the concentrations of UDP-GalNAc and UDP-GlcNAc are highly sensitive to ambient glucose levels^36^. Therefore, it is possible that the GalNAc-T enzymes as well as their donor substrate levels could critically regulate Aβ generation.

## Materials and Methods

### Materials

The sources of the materials used in this study were as follows: pCALNL5 (RDB01862, RIKEN BioResource Center), a series of GalNAcT (1, 2, 3, 4, 6, 10, 12, 13, 14, and 15)-pcDNA3.1neoGW plasmids (National Institute of Advanced Industrial Science and Technology), tissue culture medium and reagents, including Dulbecco’s modified Eagle’s medium (DMEM) from Invitrogen; protein G-Sepharose Fast Flow and immobilized streptavidin Mutein Matrix from Roche; protein molecular weight standards from Bio-Rad; BCA protein assay reagents and sulfo-NHS-LC-biotin and sulfo-NHS-SS-biotin from Thermo Fisher Scientific Inc.; and all other chemicals from Sigma or Wako Chemicals. HaloTag® Alexa Fluor 488, HaloTag®PEG biotin and HaloTag® TAMRA ligands, pFC14K HaloTag® CMV Flexi vector, and anti-HaloTag monoclonal antibody were from Promega. Other commercially available antibodies used were anti-APP (mouse, 22C11, Chemicon), anti-sAPPα (mouse, 6E10, BioLegend), anti-human APP770 (rabbit, Japan-IBL), anti-EEA1 and anti-adaptin-γ (mouse, BD Transduction Laboratories), Alexa Fluor 488-anti-Aβ (6E10, BioLegend), anti-PECAM (rat, MEC13.3, BioLegend), anti-tubulin β3 (mouse, TUJ1, BioLegend), anti-CD146 (rat, ME-9F1, BioLegend) anti-O-GlcNAc (RL2) (mouse, Thermo Fisher Scientific Inc.), anti-GAPDH (mouse, MAB374, Chemicon), anti-FLAG (M2) (mouse, Sigma), anti-ppGalNAc-T2 (rabbit, Sigma), anti-ppGalNAc-T3 (sheep, R&D Systems), anti-ppGalNAc-T6 (rabbit, Abcam), anti-Rab5 and anti-Rab7 (rabbit, Cell Signaling Technology), anti-APP (APP(C)), anti-APP OX2, and anti-human sAPPβ (rabbit, IBL-Japan).

### Construction of plasmids

APP770FLAG-pcDNA was generated as described previously^12^. For the APP770 Halo expression vector, the APP portion (forward, 5′- *gcgtacgc*ATGCTGCCCGGTTTGGCACTGCTC-3′, and reverse, 5′- *gatatc*CGTTCTGCATCTGCTCAAAGAAC-3′) was inserted into the pFC14K HaloTag® CMV Flexi vector. A series of adenovirus vectors was produced using the ViralPower Adenoviral Gateway Expression kit (Life). pENTER-FLAG APP770 was constructed by inserting the N-terminal signal peptide of APP770 (forward, 5′- *gtagac*ATGCTGCCCGGTTTGGCACTGC-3′, and reverse, 5′- *aagctt*CGCCCGACCGTCCAGGCGG-3′) and the remaining part of APP770 (forward, 5′-*tctaga*CTGGAGGTACCCACTGATG-3′, and reverse, 5′- *gcggccgc*CTAGTTCTGCATCTGCTCAAAG-3′) into the FLAG-containing pBluescript^12^, then transferred into the pENTER plasmid. The pENTER-Halo APP770 construct was created by replacing the FLAG sequence of FLAG-APP770 with the Halo portion (forward, 5′-*aagctt*GGATCCGAAATCGGTACTGGCTTTC-3′, and reverse, 5′- *tctaga*ACCGGAAATCTCCAGAGTAGACAGCC-3′). For pENTER GalNAcT constructs, the following primers were used: GalNAcT2 forward, 5′- *gcggccgc*CTAGTTCTGCATCTGCTCAAAG-3′) into the FLAG-containing pBluescript^12^, then transferred into the pENTER plasmid. The pENTER-Halo APP770 construct was created by replacing the FLAG sequence of FLAG-APP770 with the Halo portion (forward, 5′-*aagctt*GGATCCGAAATCGGTACTGGCTTTC-3′, and reverse, 5′- *tctaga*ACCGGAAATCTCCAGAGTAGACAGCC-3′). For pENTER GalNAcT constructs, the following primers were used: GalNAcT2 forward, 5′-CACCATGGCTCACCTAAAGCGACTAGTAAAA-3′ and reverse, 5′- *ggattc*ATCATTTTGGCTAAGTATCCATTTTTG-3′; and GalNAcT6 forward, 5′- CACCCTCGAGATGAGGCTCCTCCGCAGACG-3′ and reverse, 5′- GACAAAGAGCCACAACTGATGG-3′. pAd-GalNAcT2, 3, and 6 were then generated.

### Mice

All animal experiments were performed in compliance with RIKEN’s Institutional Guidelines for Animal Experiments.

### Cell culture, expression plasmids, and RNA interference

Human brain microvascular endothelial cells (BMECs, Applied Cell Biology Research Institute) were cultured in CS-C complete medium (Cell Systems) with FBS and used within four passages. Mouse primary liver sinusoidal endothelial cells^37^ and neurons^38^ were isolated and cultured as previously reported. HeLa, SK-NSH, CHO (RIKEN Cell Bank), or its mutant IdlD cells were maintained in high-glucose DMEM containing 10% FBS. For biochemical experiments, both BMECs and HeLa cells were cultured in low-glucose conditions, with MCDB131 (Sigma-Aldrich) and DMEM containing 10% FBS, respectively, for at least 24 h. The BMECs were then transfected using Nucleofector (Lonza, Basi Nucleofector Kit for primary endothelial cells, program M003), and the HeLa cells were transfected using FuGENE6 reagent (Promega). For knockdown experiments, Stealth RNAis (Invitrogen) were used. HeLa cells at 50% confluency on 6-cm dishes were infected with hAPP770FLAG-pAd. After 24 h, the cells were transfected with 50 pmol control siRNA (Stealth RNAi negative control medium GC Duplex) or siRNA for GalNAcT2 (HSS103983), GalNAcT3 (HSS103984), or GalNAcT6 (HSS117436) using Lipofectamine RNAiMAX Transfection Reagent (Thermo Fisher Scientific). After 16 h, the culture medium was changed to low-glucose DMEM medium containing 2% FBS. After 24 h, the cells and medium were collected for further analysis.

### Human samples

The clinical study was approved by the ethical committees of RIKEN, Tokyo Metropolitan Institute of Gerontology, Tokyo Metropolitan Geriatric Hospital, and Fukushima Medical University. Frozen tissues from postmortem brain were obtained from the Brain Bank for Aging Research, which consists of samples from consecutive autopsy cases from a general geriatric hospital with informed consent obtained from the relatives for each autopsy. The handling of the brain tissue and the diagnostic criteria have been described previously^39^. Cerebrospinal fluid (CSF) samples were collected from patients with Alzheimer’s disease.

### Real-time PCR

Total RNA from cultured cells was extracted using TRIzol (Invitrogen). One microgram of total RNA was reverse-transcribed using the SuperScript III First-Strand Synthesis System (Invitrogen) with random hexamers. The cDNAs were mixed with TaqMan Universal PCR master mix (Life Technologies) and amplified using an ABI PRISM 7900HT sequence detection system (Applied Biosystems). All primers and probes, *GALNT2* (Hs00189537_m1), *GALNT3* (Hs00237084_m1), *GALNT6* (Hs00926629_m1), and *18S rRNA* (Hs99999901_s1), were from Applied Biosystems. The levels of mRNA were normalized to the corresponding ribosomal RNA levels.

### Immunofluorescence

Cells metabolically labeled with Ac_4_GalNAz were fixed with 4% paraformaldehyde in PBS, treated with 0.25% Triton X-100 for 15 min, and labeled with Alexa555-alkyne, in accordance with the manufacturer’s instructions (Invitrogen/Click-iT Cell Reaction Buffer Kit). For the analysis of Halo-APP770 uptake, the cells were incubated with non-permeable HaloTag® Alexa 488 ligand or biotin ligand, or permeable HaloTag® TAMRA ligand for 5–30 min before fixation. For double staining, the cells were incubated at 4°C with antibodies to the following: APP OX2 (1:75), adaptin-γ (1:100), EEA1 (1:100), Rab5 (1:150), Rab7 (1:100), GANLT2 (1:250), and DAPI. As antibodies for endogenous Rab11 were ineffective for immunocytochemistry, FLAG-tagged Rab11 was expressed in BMECs, and detected with anti-FLAG (M2, 1:100) to obtain a clear Rab11 signal. The next day, the cells were washed three times and incubated with the appropriate fluorescently labeled goat secondary antibodies (1:1000, Invitrogen) for 1 h at room temperature. Images were taken using an Olympus FV-1000 confocal microscope, with data acquisition and quantification of the signal or co-localization area being carried out using FV10-ASW ver.1.7 software (Olympus).

### Isolation of endothelial cells

Endothelial cells were isolated from mouse brains as previously reported^12, 40^, except for the use of anti-CD146 antibody coupled with Dynabeads sheep anti-Rat IgG (Thermo Fisher Scientific).

### Western and lectin blot

The samples were subjected to SDS-PAGE (5%–20% gradient gel) and transferred to nitrocellulose membranes. For western blot analyses, following incubation with 5% non-fat dried milk in TBS-containing 0.1% Tween-20, the membranes were incubated with anti-APP 22C11 (1:1000 dilution), anti-APP(C) (1:1000 dilution), anti-O-GlcNAc RL2 (1:500 dilution), or anti-actin (1:500 dilution) antibodies. Appropriate horseradish peroxidase-conjugated donkey anti-goat IgG (Jackson ImmunoResearch Laboratories) or anti-mouse and anti-rabbit IgG (GE Healthcare) antibodies were used as the secondary antibodies (1:1000 dilution). For the lectin pull-down experiment, the lysates (50 μg of protein) were incubated with 20 μl of *Arachis hypogaea* (PNA)-coupled agarose (J-oil Mills), and subjected to western blot analysis with anti-APP(C). Signals were detected with SuperSignal West Dura Extended Duration Substrate (Thermo Fisher Scientific) using ImageQuant LAS-4000mini (GE Healthcare). The intensity of the resultant protein bands was quantified using ImageQuant TL software (GE Healthcare).

### Azide sugar labeling

BMECs were metabolically labeled with 100 μM Ac_4_GalNAz or Ac_4_MaNAz for 6 h. The Click-iT protein reaction buffer kit with Alexa555-labeled alkyne was used to label the fixed cells. For biochemical detection, the cell lysates (350 μg of protein) were incubated with TAMRA-conjugated DIBO for 3 h. sAPP in the medium was pulled down with heparin-agarose (Thermo Fisher Scientific Inc.). The samples were then subjected to SDS-PAGE analysis. Fluorescent signals on the gel were visualized with Typhoon 9400 (GE Healthcare).

### Glycosidase treatment

Glycoproteins in the cell lysates (20 μg of protein) or medium that were pulled down with heparin-agarose (0.5 ml) were denatured and incubated with *Arthrobacter ureafaciens* sialidase (Nacalai Tesque, 4 milliunits) and/or O-glycosidase (New England BioLabs, 80,000 units), or peptide N-glycanase (PNGase, New England Biolabs, 1,000 units) for 6 h. To remove O-GlcNAc, cell lysates (100 μg of protein) were incubated with O-GlcNAcase (5 μg, R&D Systems) at 37°C for 2 h.

### Cell surface biotinylation assay

To measure the half-life of cell surface APP, or for biochemical analysis of cell surface biotinylated APP, Sulfo-NHS-LC-biotin was used from BMECs for 30 min at 4°C. After washing the plates three times with 0.1 M glycine in PBS (pH 8.0) and once with PBS alone, cell lysates were prepared with TPER buffer (Thermo Scientific). For the internalization assay, BMECs were labeled with NHS-SS-biotin, and cultured for 0, 5, 10, or 60 min, after which cell surface biotin was removed using glutathione reagent as previously described^12, 41^. Biotinylated proteins were pulled down with immobilized streptavidin. To check the incorporation of azide sugars in the biotinylated APP, following cell surface biotinylation with sulfo-NHS-LS-biotin, BMECs expressing APP770FLAG or control vector were metabolically labeled with azide sugars and lysed. Biotinylated proteins bound to immobilized streptavidin were eluted with 2 mM biotin, after which APP770FLAG was immunoprecipitated with anti-FLAG M2-agarose.

### *In vitro* GalNAcT assay

Human APP770-derived peptides, APP-OX2 (QSLLKTTQEPLA), APP^361-372^ (PVKLPTTAASTP), and APP^646-657^ (ADRGLTTRPGSG), which were tagged with a 9-fluorenylmethyloxycarbonyl group at their N-terminus, were used as acceptor substrates. A series of recombinant soluble FLAG-GalNAcTs was purified from the medium of overexpressing COS cells by immuno-affinity chromatography using M2 agarose. The concentration of each GalNAcT-FLAG enzyme was measured by performing immunoblot analysis together with standard FLAG-BAP fusion protein (Sigma-Aldrich) using the M2 antibody, and adjusted. The standard enzyme reaction mixture containing 15 μM acceptor substrate, and the purified enzyme, 0.5 mM UDP-GalNAc in 25 mM Tris-HCl (pH 7.4), 5 mM MnCl_2_, and 0.1% Triton X-100 in a final volume of 20 μl, was incubated at 37 °C for 16 h, after which the reaction was terminated by boiling^28^. The reaction products were purified with Millipore Ziptips and mixed with a matrix (2,5-dihydroxybenzoic acid), which was analyzed by Bruker Ultraflex MALDI mass spectrometry in the positive ion mode.

### Phylogenic analysis

Amino acid sequences of human ppGalNAc-T were obtained from the RefSeq database^42^. Evolutionary analysis was conducted in MEGA6^43^ using the UPGMA method.

### Quantification of sAPP and Aβ

HeLa cells and BMECs were infected with adenovirus preparations for APP770-FLAG overexpression. HeLa cells and primary neurons were transfected with GalNAcT2-, T3-, or T6-encoding pcDNA using FuGENE6 and Lipofectamine 2000 (Invitrogen), respectively. BMECs were infected with adenovirus to express GalNAcT2, T3, or T6. To measure sAPP770, sAPP770β, Aβ40, and Aβ42 in the medium, a Human sAPP Total Assay Kit, an sAPPβ-w Assay Kit (highly sensitive) (IBL-Japan), and human amyloid β (1–40) and (1–42) assay kits were used, respectively^46^. To measure O-GalNAc glycans on APP770, a 96-well plate coated with anti-OX2 antibody (IBL-Japan) was incubated with BMEC lysate for 16 h. *Sambucus sieboldiana* agglutinin (SSA)-biotin (J-oil Mills, 1:1000) and streptavidin-HRP (1:1000) were then used for detection.

### Determination of O-glycosylation sites in APP770

HA-tagged human APP770 was expressed in HEK293T cells, and HA-sAPP770 purified from culture medium was precipitated in acetone at −30°C for 16 h and then centrifuged at 12,000 × g for 10 min. The precipitated sample was reduced in 10 mM dithiothreitol (DTT) for 30 min at 56°C, and alkylated with 20 mM iodoacetamide for 40 min at 25°C in the dark. Then, the proteins were digested with 1.5 µg of Trypsin/Lys-C mix (Promega) for 16 h at 37°C (800 rpm). The O-glycopeptides were precipitated with a five-fold volume of ice-cold acetone by centrifugation at 12,000 × g for 10 min (Fraction 1). The supernatant was collected in a separate tube and dried in a speed vac. concentrator. O-glycopeptide from the supernatant was enriched with GlycOCATCH (Genovis), in accordance with the manufacturer’s instructions. In brief, the supernatant reconstituted in 0.1% Triton x-100 in TBS was reacted with GlycOCATCH affinity resin at room temperature for 2 h with 10 units of SialEXO. The resin was washed three times with 0.5 M sodium chloride in TBS, and then eluted by incubating with 8 M urea for 5 min at room temperature with mixing (Fraction 2). Both fractions were combined, purified by GL-Tip SDB (GL Sciences), and then subjected to LC/MS. To identify the O-glycosylation site, a portion of the glycopeptide fraction was treated with OpeRATOR (Genovis) at 37°C overnight.

The O-glycopeptides were separated on an EASY-nLC 1000 (Thermo Fisher Scientific) with an Acclaim PepMap100 C18 LC column (75 µm×20 mm, 3 µm; Thermo Fisher Scientific) and a Nano HPLC Capillary Column (75 µm×120 mm, 3 µm, C18; Nikkyo Technos). The eluents consisted of water containing 0.1% v/v formic acid (pump A) and acetonitrile containing 0.1% v/v formic acid (pump B). The O-glycopeptides were eluted at a flow rate of 0.3 µL/min with a linear gradient from 0% to 35% B over 40 min. Mass spectra were acquired on a Q Exactive mass spectrometer (Thermo Fisher Scientific) equipped with Nanospray Flex Ion Source (Thermo Fisher Scientific) operated in the positive ion mode. We used an Xcalibur 4.4 workstation (Thermo Fisher Scientific) for MS control and data acquisition. The spray voltage was set at 1.8 kV, while the capillary temperature was kept at 250°C. The full mass spectra were acquired using an *m/z* range of 350–2000 with a resolution of 70,000. The product ion mass spectra were acquired against the 10 most intense ions using a data-dependent acquisition method with a resolution of 17,500 with normalized collision energy (NCE) of 27.

### Analysis of GalNAz-incorporated APP

FLAG-APP770 was expressed in BMECs using an adenoviral system and purified from cell lysates with anti-FLAG M2-agarose (Sigma-Aldrich). The lyophilized sample (30 μg of protein) was reduced with dithiothreitol (10 mg, 50°C for 1 h) and alkylated with iodoacetamide (20 mg, room temperature for 30 min in the dark). After the reaction mixture had passed through a Nap-5 column (GE Healthcare) to remove excess dithiothreitol and iodoacetamide, the sample was digested with trypsin (2 μg, Promega) in 50 mM ammonium bicarbonate (100 μl) for 16 h at 37°C. After boiling for 10 min, the sample was evaporated to dryness. The dried residue was dissolved with 12 μl of mobile phase (A), and a portion of it (5 μl) was used for LC-ESI MS and MS/MS analyses to determine the presence of glycopeptide containing GalNAz. The glycopeptide mixtures were separated using an ODS column (Develosil 300ODS-HG-5, 150 × 1.0 mm i.d.; Nomura Chemical). The mobile phases were (A) 0.08% formic acid and (B) 0.15% formic acid/80% acetonitrile. The column was eluted with solvent A for 5 min, at which point the concentration of solvent B was increased to 40% over 55 min at a flow rate of 50 μl/min using an Accela HPLC system (Thermo Fisher Scientific). The eluate was continuously introduced into an ESI source (LTQ Orbitrap XL; Thermo Fisher Scientific at the Natural Science Center for Basic Research and Development, Hiroshima University). MS and MS/MS spectra were obtained in the positive ion mode using Orbitrap (mass range: *m/z* 300 to 3000) and Iontrap (data-dependent scan of the top 3 peaks from a prepared list using CID), respectively. The voltage of the capillary source was set at 4.5 kV and the temperature of the transfer capillary was maintained at 300°C. The capillary voltage and tube lens voltage were set at 15 V and 50 V, respectively.

### Data availability

This study includes no data deposited in external repositories.

## Acknowledgments

We thank Dr. Monty Krieger (Massachusetts Institute of Technology) for the CHO IdlD cells, Dr. Takeshi Ijuin (Kobe University) for the p3xFLAG-CMV8 mRab11 vector, Dr. Izumu Saito (University of Tokyo) for pCALNL5, and Dr. Hisashi Narimatsu (National Institute of Advanced Industrial Science and Technology) for the series of ppGalNAc-Ts in pcDNA3.1neoGW or pFLAG-CMV3-DEST plasmids. We also thank R. Hayashi and T. Chiaki from Leica Microsystems for their technical help in obtaining super-resolution images. We additionally thank T. Tajima (BSI-Olympus Collaboration Center, Brain Science Institute) for his technical support in analyzing super-resolution imaging. Moreover, we wish to express our appreciation to Dr. Toshiyuki Yamaji (National Institute of Infectious Diseases) for providing critical comments on our manuscript. This work was supported by a grant to RDC (P41GM103694), the Mitsubishi Foundation to SK, and by the Japan Society for the Promotion of Science (JSPS) (Grant numbers 26117522 and 25430122 to SK and 25840043 to YT).

## Author contributions

S.K. conceived the study and wrote the manuscripts. Y.T. and J.I. mainly designed and performed the experiments. K.T. and H.S. performed additional experiments. M. N., D.T., and N.K. contributed to MS analysis. Y.Y. performed bioinformatics analysis. K.T. prepared click chemical probe. Y.S., H.M., and T.E., contributed to analyze human samples.

## Conflict of interest

The authors declare no competing financial interests.

The Paper Explained

Problem

Aβ deposition in brain parenchyma and blood vessels is a critical pathological hallmark in Alzheimer’s disease (AD). Endothelial amyloid β precursor protein (APP) and neuronal APP contribute to the disease pathogenesis. APP has two N-glycans and multiple O-glycans, but how such glycosylation affects intracellular APP trafficking and Aβ generation is not fully understood.

Results

We analyzed APP glycosylation and intracellular trafficking in neuronal and endothelial cells, and found a significant difference. Neuronal APP follows the typical glycosylation pathway, while we unexpectedly observed that, in endothelial cells, APP lacking O-glycans, but having N-glycans, is externalized to the endothelial cell surface and transported back to the Golgi apparatus, where it then acquires O-glycans. Knockdown of the genes encoding enzymes initiating APP O-glycosylation significantly reduced Αβ production.

Impact

Our finding raises the possibility that a non-classical glycosylation pathway contributes to Aβ generation and can be considered a novel therapeutic target.

## Expanded View Figure

**Figure EV1.**
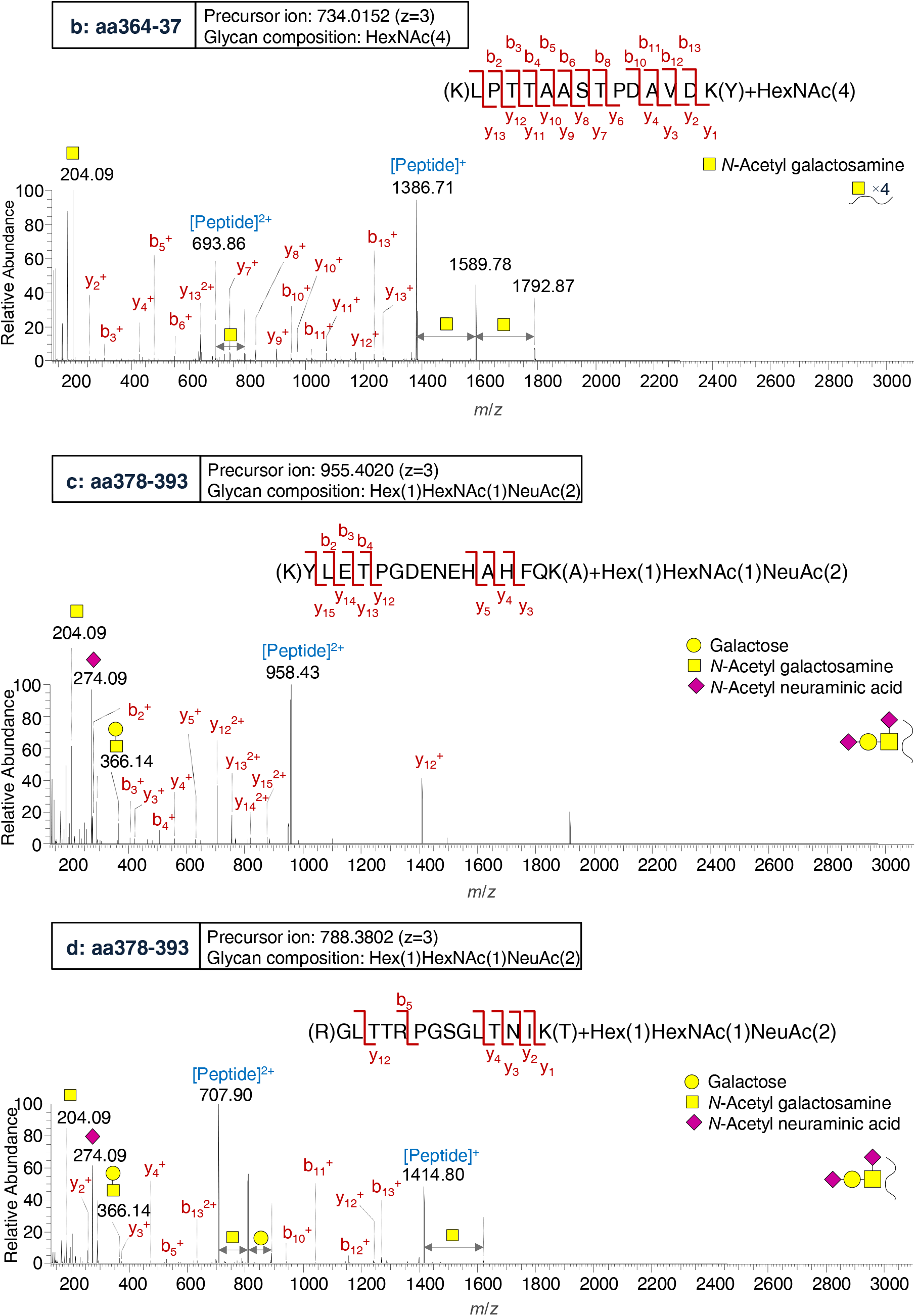
Representative product ion spectra of APP770 O-glycopeptide. aa364–377 arising from the precursor ion at *m/z* 734.0152 (z=3), aa378–393 arising from the precursor ion at *m/z* 955.4020 (z=3), and aa649–662 arising from the precursor ion at *m/z* 788.3802 (z=3).

**Figure EV2.**
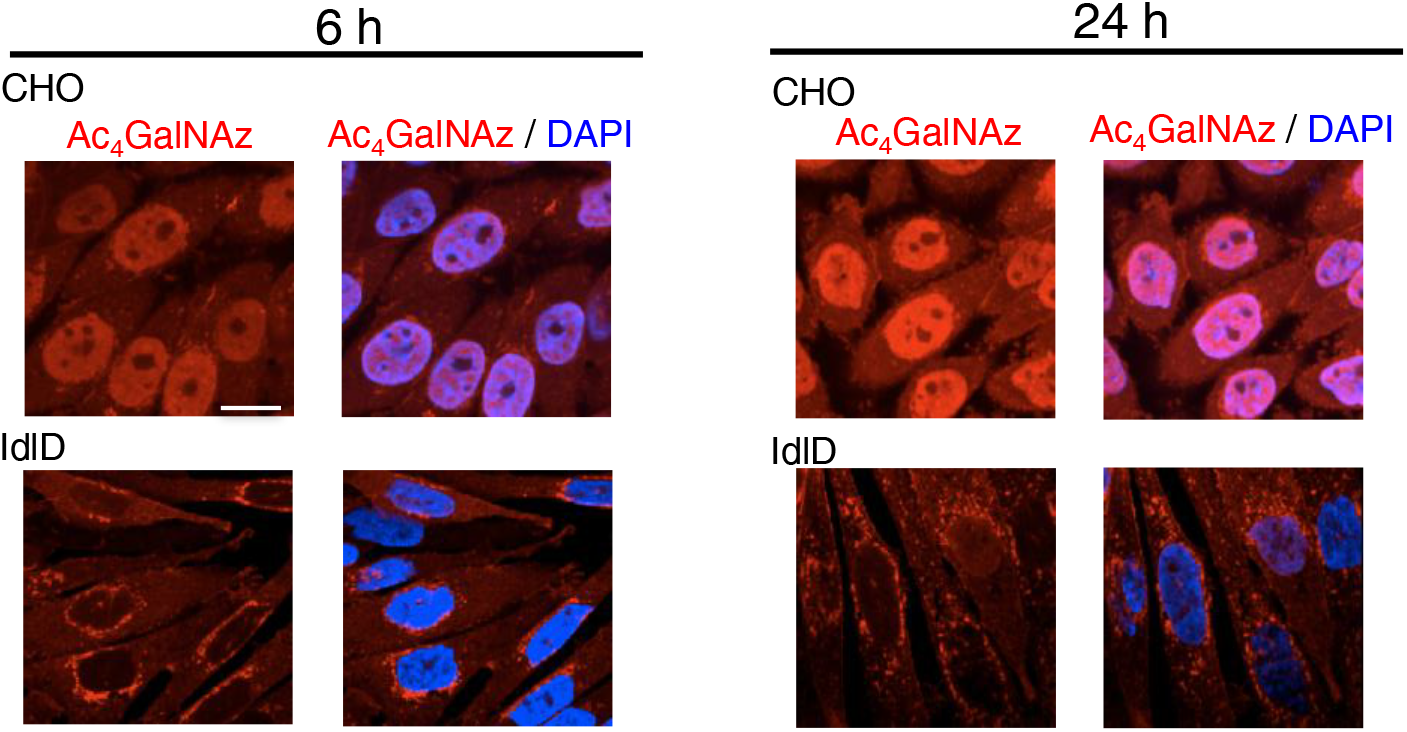
Metabolic labeling with GalNAz results in nuclear azide signals derived from O-GlcNAc signals. IdlD cells, which lack UDP-galactose epimerase (GALE) activity, and their parental CHO cells were incubated with GalNAz. After 6 or 24 h of incubation, the cells were fixed, reacted with Alexa555-alkyne and DAPI, and analyzed by immunofluorescence microscopy. Scale bar, 20 μm.

**Figure EV3.**
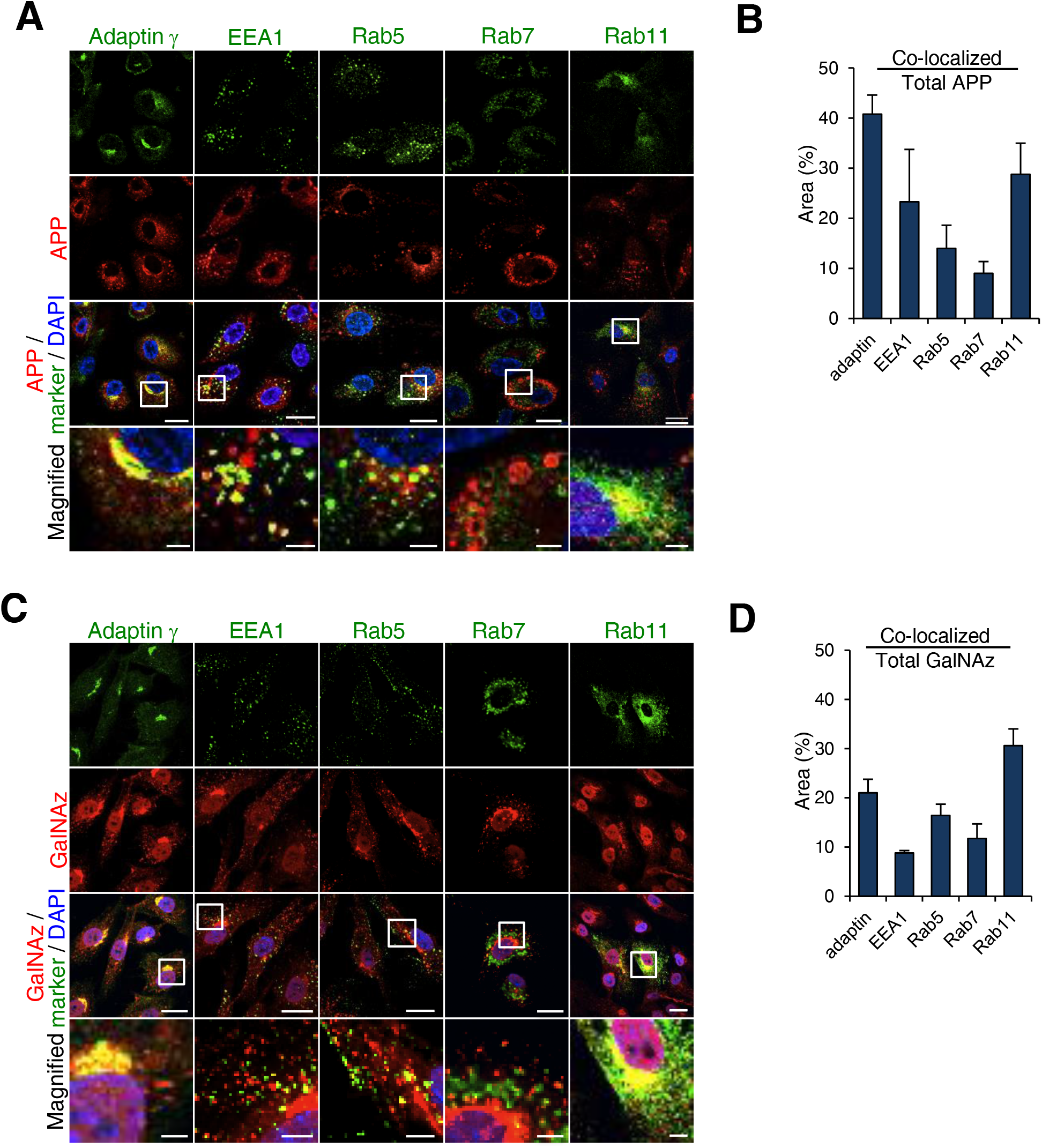
O-GalNAz glycans are enriched in the recycling endosome. A Immunostaining of BMECs with APP-FLAG and several organelle markers (adaptin-γ, EEA1, Rab5, Rab7, and Rab11). Scale bar, 20 μm (and 4 μm for magnified image). B Quantitative analysis of the percentage of APP and organelle marker co-localization is shown as mean ± SEM. (n=4) in (A). C Immunostaining of BMECs for O-GalNAz glycans and several organelle markers. Scale bar, 20 μm (and 4 μm for magnified image). D Quantitative analysis of the percentage of O-GalNAz glycan and organelle marker co-localization is shown as mean ± SEM (n=4) in (C).

**Figure EV4.**
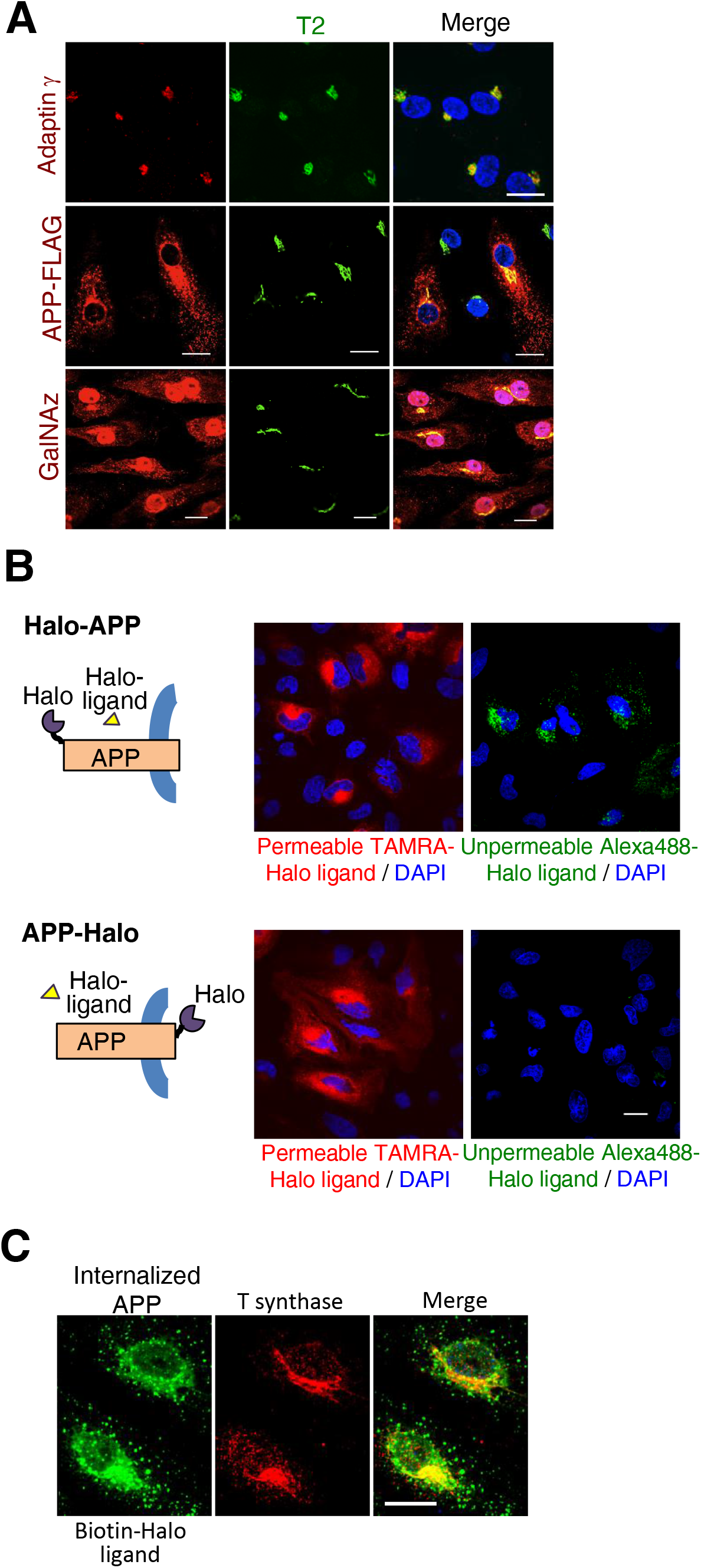
O-glycosylation enzymes co-localize with GalNAz and internalized APP. A BMECs expressing APP-FLAG were cultured with GalNAz for 6 h. Following fixation, the cells were reacted with Alexa555-labeled alkyne. Immunostaining analysis of GalNAc-T2 was performed for adaptin-γ, APP-FLAG, and GalNAz. Scale bar, 20 μm. B HeLa cells expressing Halo-APP or APP-Halo were reacted with the non-permeable HaloTag® Alexa488 ligand or the permeable HaloTag® TAMRA ligand. Scale bar, 20 μm. C BMECs expressing Halo-APP in an adenovirus system were incubated with HaloTag® biotin ligand for 30 min. After fixation, the cells were reacted with SA-Alexa488 and then immunostained for T-synthase. Scale bar, 20 μm.

**Figure EV5.**
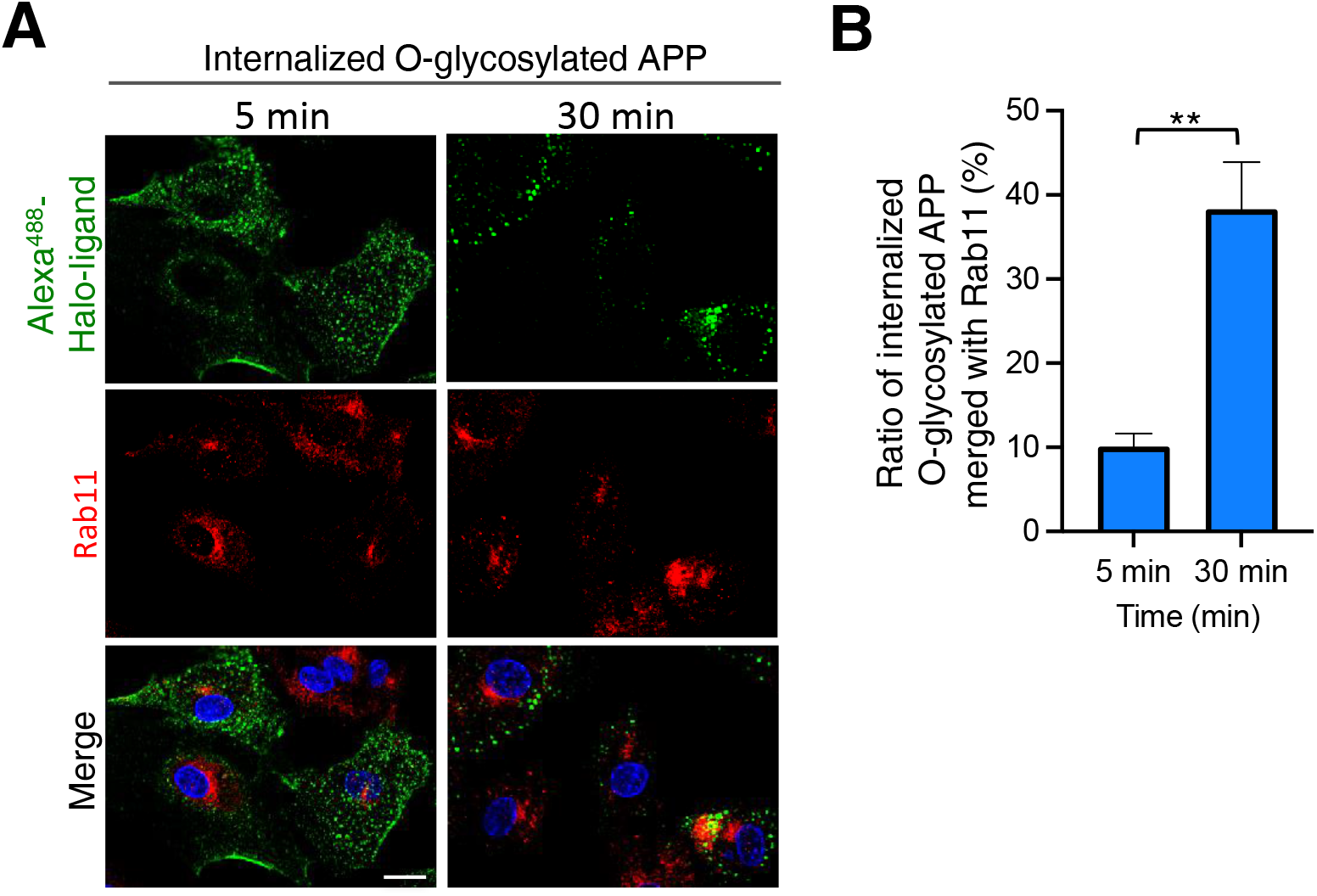
Maturation of internalized O-glycosylated APP vesicles upon internalization. A The cells were fixed and immunostained for Rab11. Scale bar, 20 μm. B The rate of co-localization of internalized O-glycosylated Halo-APP with Rab11 after 5 or 30 min of Halo-ligand uptake was determined as mean ± SEM (n=5).

---

**Appendix Figure S1.**
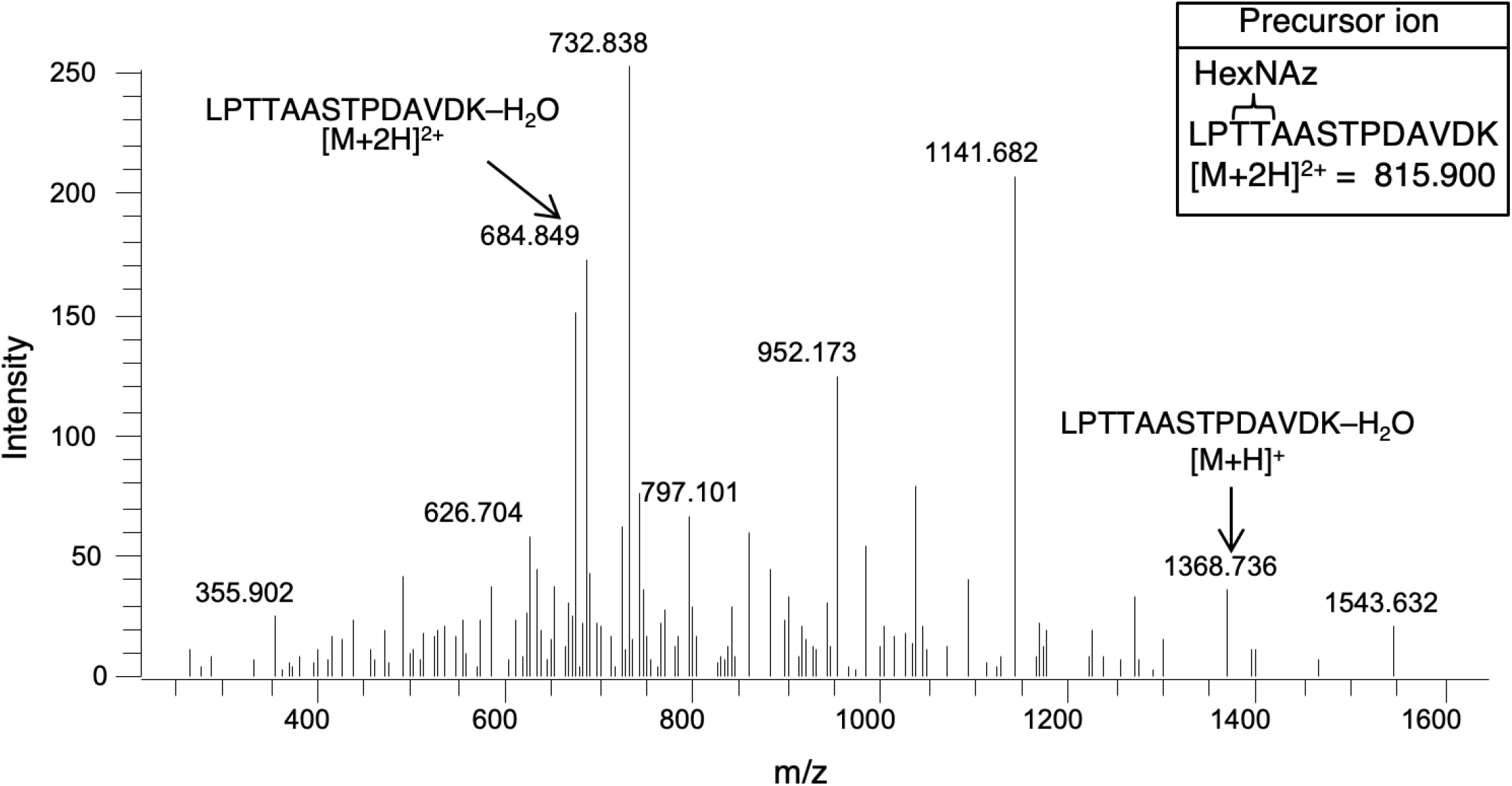
Incorporation of GalNAc as part of O-glycans in APP770. APP-FLAG, expressed using an adenovirus system in BMECs metabolically labeled with GalNAz, was immunopurified with an M2-coupled agarose column. The MS/MS of a precursor ion at *m/z* 815.90 representing [M+2H]^2+^ of LPTTAASTPDAVDK+HexNAz gives two indicative fragment ions at *m/z* 1368.74 and *m/z* 684.85 representing [M+H]^+^ and [M+2H]^2+^ of LPTTAASTPDAVDK-H_2_O, respectively. These fragment ions show the presence of GalNAz in the peptide LPTTAASTPDAVDK.

**Appendix Figure S2.**
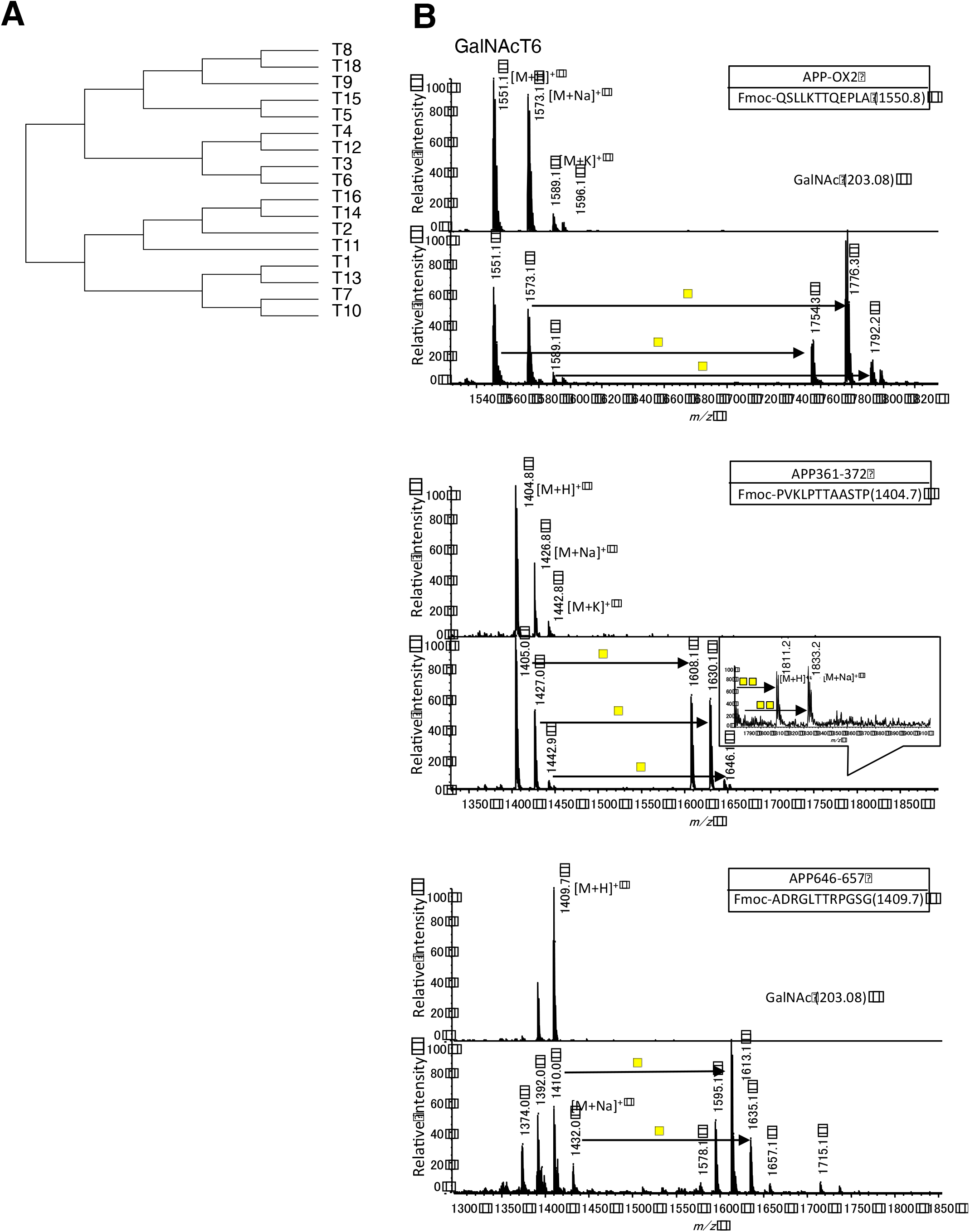
*In vitro* GalNAc-T assay shows the incorporation of GalNAc into APP-derived peptide. A. Phylogenic analysis of the human GalNAc-T family. B. Soluble recombinant GalNAc-T6 was incubated with three kinds of APP-derived peptide and UDP-GalNAc. The reaction products were partially purified by Millipore Ziptips and analyzed by mass spectrometry (AXIMA-QIT TOF-MS, Shimadzu Biotech). MS spectra of the reaction products (lower) show the incorporation of one or two GalNAc residues into the peptide substrate (upper).

